# Heightened gaze responsiveness and mutual gaze instability at close distances characterize dyadic coordination in a marmoset model of autism

**DOI:** 10.1101/2025.11.17.688777

**Authors:** Madoka Nakamura, Tomoko Sato, Tohru Kurotani, Kazuhisa Sakai, Anna Maria Hadjiev, Shuntaro Sasai, Koki Mimura, Nobuyuki Kawai, Tsuyoshi Shimmura, Noritaka Ichinohe

## Abstract

Autism spectrum disorder (ASD) is associated with differences in social interaction, including alterations in gaze behavior and interpersonal distance. Although most assessments of ASD remain individual-centric, emerging human studies indicate that social difficulties may arise from atypical dyadic coordination rather than from reduced social interest alone. We examined dyadic coordination in a non-human primate model of ASD, focusing on gaze and inter-individual distance during minimally constrained first encounters between prenatal valproic acid-exposed marmosets (VPA) and unexposed conspecifics (UE). Markerless pose estimation quantified frame-by-frame gaze direction and spacing, comparing VPA-UE with UE-UE dyads. VPA-UE pairs exhibited a higher proportion of mutual-gaze events than UE-UE pairs. VPA individuals showed more rapid looks in response to the partner’s gaze, yet spent less time looking at the partner when unobserved. VPA-UE dyads maintained shorter inter-individual distances than UE-UE dyads, and closer spacing was associated with decreased stability of mutual gaze. These findings reveal a dyadic coordination pattern in VPA-UE pairs, with rapid responsiveness coupled with distance-dependent fragility, paralleling features reported in human ASD. Our dyad-centric, markerless framework yields ecologically grounded readouts and potential intervention targets that leverage intact social responsiveness while accommodating altered spatial coordination.

## Introduction

Autism spectrum disorder (ASD) is often characterized by atypical regulation of gaze and interpersonal distance, two core channels of social coordination^1–5^. However, a growing literature indicates that these differences do not simply reflect reduced social motivation. Empirical work shows that many autistic individuals orient rapidly, sometimes reflexively, to others’ eyes but then avert their gaze as proximity heightens arousal^6–8^, whereas others wish to remain close yet still step back when a partner approaches^9,10^. These context-dependent patterns point to difficulties in regulating arousal and proximity rather than diminished social interest, with gaze and distance operating as coupled channels for modulating the intensity of social engagement.

This view aligns with dyadic frameworks that conceive social behavior as emerging from real-time coordination between partners. The “double empathy problem”^11^ posits that difficulties arise when two individuals bring different sensory thresholds, expectations, or pacing styles into the same encounter^12,13^. Hyperscanning studies support this, showing that neurotypical partners interacting with autistic individuals recruit additional prefrontal and occipital resources, providing evidence that the challenge is not solely within the autistic individual but within the mismatched dyad^14^. From this perspective, the timing of gaze initiation, the strength of responsiveness to a partner’s look, and the stability of mutual engagement reflect ongoing adjustments to spatial proximity and arousal. These accounts therefore predict that ASD-related differences will be expressed not only in mean levels of gaze or distance, but in the moment-to-moment contingencies linking one partner’s actions to the other’s reactions.

However, most human experimental paradigms rely on screen-based, movement-restricted, single-participant designs. Interpersonal distance is held constant, and eye contact is sampled outside the context in which it naturally occurs. These constraints limit our ability to fully understand how gaze and distance jointly regulate the dynamics of social interaction^15–17^.

A species that naturally integrates direct gaze with fine-grained spatial negotiation is therefore particularly valuable for this problem. The common marmoset (*Callithrix jacchus*) combines several traits that are directly relevant to human face-to-face interaction: it readily tolerates neutral direct gaze, engages in brief bouts of mutual gaze during affiliative encounters, and finely adjusts body orientation and spacing in response to a partner’s movements^18–24^. This reliance on visual social monitoring, in contrast to the more olfactory-focused communication of rodents and the more threat-related interpretation of direct gaze in macaques, makes marmosets particularly suitable for probing how gaze and distance are co-regulated in primate social interaction.

Prenatal exposure to valproic acid (VPA) yields a marmoset model that recapitulates multiple ASD-like features within this gaze–distance coordination system, including reduced affiliative calling^25,26^, difficulty sustaining stable mutual engagement^27,28^ and heightened vigilance^29^. These traits instantiate an approach-avoidance tension closely aligned with accounts of ASD in humans, but expressed in a species whose native social repertoire preserves the fine-grained interplay between gaze, distance, and arousal.

Here we analyzed minimally constrained first-encounter interactions between VPA-exposed and unexposed (UE) marmosets using frame-by-frame pose estimation to quantify how gaze and inter-individual distance are coordinated in real time. The central question was whether ASD-like differences would manifest not as a uniform reduction in social engagement, but as a distinct constellation of heightened reactivity, rapid yet fragile formation of mutual gaze, and distance-dependent breakdowns of coordination, patterns that are predicted by dyadic accounts of ASD but largely invisible in single-participant paradigms. This approach enabled us to evaluate both short-timescale responsiveness and longer-timescale spatial positioning within the same interaction, describing social difficulties not as uniform deficits but as distance-dependent alterations in dyadic coordination.

## Results

We investigated whether prenatal VPA-exposed ASD model marmosets display distinctive gaze patterns when encountering an unfamiliar conspecific for the first time, and further examined whether their typically developing partners also exhibited altered gaze patterns in this interactive context. In a first-encounter test with a transparent barrier allowing visual and auditory contact but preventing physical interaction (Fig. 1a), either a VPA-exposed or an unexposed (UE) marmoset (*n* = 10 per group; Supplementary Table S1) was placed in a rectangular arena as the target individual and paired with an unfamiliar, same-sex UE partner confined in a circular container positioned to the left or right of the target arena (Fig. 1a). Head positions (forehead, left ear, and right ear) were tracked frame by frame using DLC (Fig. 1b; see Supplementary Video S1 for a representative example), and the gaze angle (*θ*) was defined relative to the vector pointing toward the partner’s forehead (Fig. 1c).

**Fig 1.**
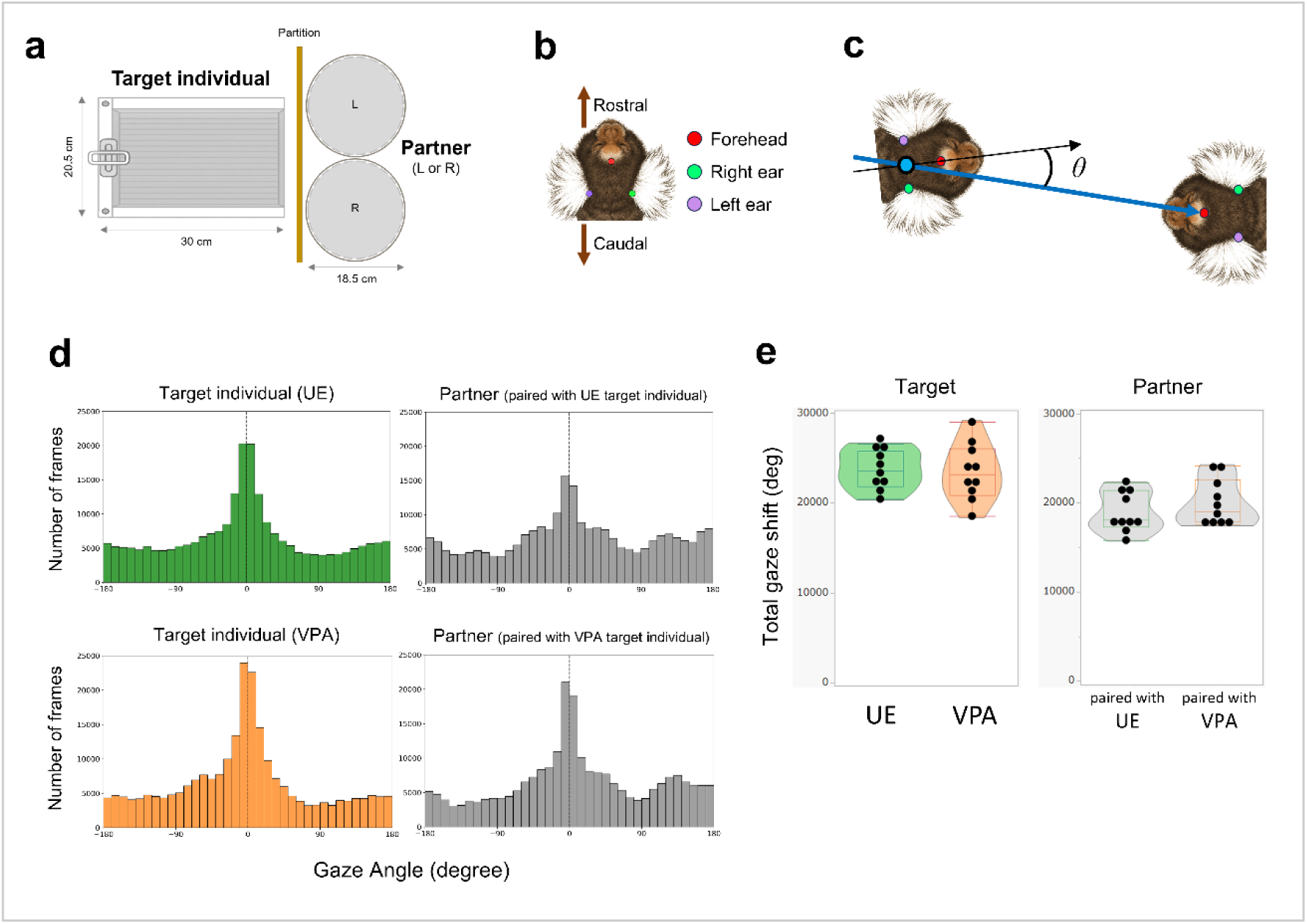
Experimental setup and quantification of gaze angle and gaze shifts. (**a**) Overhead view of the experimental apparatus. The target individual (either VPA or UE) was placed in a rectangular arena, and an unfamiliar partner was placed in a circular container positioned to the left or right of the target arena. The partner’s position was counterbalanced across trials. The partition between the two individuals was lifted upward by the experimenter at the start of the experiment, enabling them to see each other. (**b**) Head orientations of both subjects were tracked using DLC: forehead (red), right ear (green), and left ear (purple). Rostral-caudal orientation is indicated by arrows. (**c**) The gaze angle (*θ*) was calculated as the angle between two vectors: one from the midpoint of the left and right ears to the forehead of the subject, and the other from the midpoint of the left and right ears to the forehead of the other conspecific. (**d**) Frequency (number of frames) distributions of gaze angles (*θ*) for all target individuals and their partners in the two groups. (**e**) Total gaze shift (deg) for the target individuals (left) and their partners (right), defined as the trial-wise sum of absolute frame-to-frame changes in gaze angle.

Across groups, the frame-wise frequency distributions of gaze angles showed a pronounced peak at 0°, indicating that looking directly at conspecifics was the most common orientation during the trial (one-sample V tests against 0° on pooled frame-wise data, all *p* < 0.0001; Watson’s U^2^ values ≈ 370–1587; Fig. 1d). Total gaze shift, defined as the cumulative sum of absolute frame-to-frame changes in gaze angle over the trial, did not differ significantly between groups, indicating that the way subjects oriented and redirected their gaze during the trial did not differ between groups (*linear mixed model*, LMM: *β* = −299.8 ± 1208.8, *t*(18) = −0.248, *p* = 0.807 for target individuals, Mann-Whitney U test: *U* = 117, *p* = 0.385 for partners; Fig. 1e).

### VPA**–**UE pairs exhibit an elevated proportion of mutual gaze compared to UE**–**UE pairs

When defining “looking at the conspecific” as a gaze angle (*θ*) within ±30°, the proportion of time spent looking at the partner did not differ significantly between VPA and UE target individuals (LMM: *β* = 4.76 ± 3.75, *t*(18) = 1.27, *p* = 0.22; Fig. 2a). Among partners, the proportion was slightly higher in those paired with VPA target individuals, but the difference was not statistically significant (Mann-Whitney U test: *U* = 126, *p* = 0.121; Fig. 2a). However, VPA target individuals exhibited more looking events than UE target individuals (LMM: *β* = 12.27 ± 5.39, *t(*18) = 2.28, \**p* = 0.0353; Fig. 2b). Interestingly, partners also looked more frequently when paired with a VPA target than with a UE target (Mann-Whitney U test: *U* = 134.5, \**p* = 0.0282; Fig. 2b). This partner-side change parallels human hyperscanning findings^14^, in which neurotypical individuals also show altered responses when interacting in real time with autistic partners.

**Fig 2.**
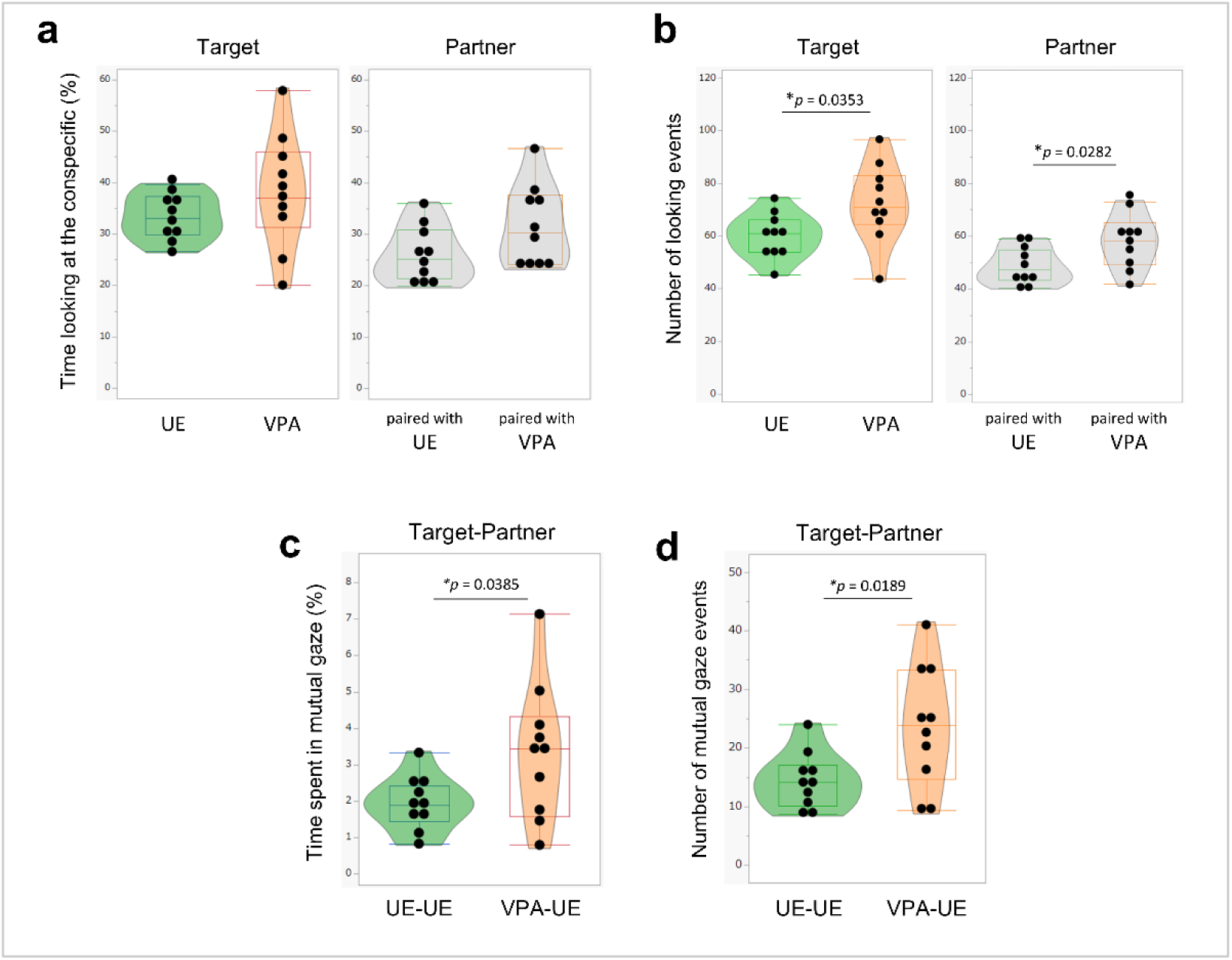
VPA–UE marmoset pairs show increased gaze events and mutual gaze compared with UE–UE pairs. (**a**) Proportion of time spent looking at the partner (gaze angle within ±30°) in target individuals (left) and their partners (right). No significant differences were found between VPA and UE groups (*p* = 0.298) and between the partners (*p* = 0.121). (**b**) Number of looking events during dyadic interaction. Target VPA individuals exhibited more looking events compared with UE controls (\**p* = 0.0353), and their partners also looked more frequently when paired with a VPA target than with a UE target (\**p* = 0.0282). (**c**) Proportion of mutual gaze (both individuals having gaze angles within ±10°) in UE–UE and VPA–UE pairs. VPA–UE pairs showed significantly higher mutual gaze (\**p* = 0.0385). (**d**) Number of mutual gaze events was significantly higher in VPA–UE pairs (\**p* = 0.0189). Statistical differences were assessed using a linear mixed model (LMM) for repeated-measures data obtained from the target individuals and the target-partner pairs, and the Mann-Whitney U test for independent measurements from the partners. * = *p* < 0.05.

When “mutual gaze” was defined as both individuals simultaneously having gaze angles within ±10°, a more stringent criterion intended to isolate face-to-face orientation, VPA–UE pairs exhibited a significantly higher proportion of mutual gaze than UE–UE pairs (LMM: *β* = 1.42 ± 0.63, *t*(18) = 2.23, \**p* = 0.0385; Fig. 2c). Consistently, the number of mutual gaze events was also significantly greater in VPA–UE pairs compared with the controls (LMM: *β* = 9.27 ± 3.59, *t*(18) = 2.58, \**p* = 0.0189; Fig. 2d). These group-level differences were also apparent at the level of single sessions; representative dyadic gaze timelines from each pair are shown in Supplementary information (Supplementary Fig. S1). To characterize how mutual gaze ceased, we examined whether the target or the partner tended to terminate it. In both pair types, target individuals ended mutual gaze more often than partners, and the rate for target individuals was comparable between VPA and UE groups (Supplementary Fig. S2).

### VPA marmosets exhibit fewer one-sided looks and heightened gaze responses

To further investigate the fine-grained dynamics of interactive gaze, we focused on the reciprocal relationship of looking and being looked at between target subjects and their partners (see Methods). One-sided looks were computed for each individual as the fraction of its looking events for which the other individual did not simultaneously orient toward it. The proportion of one-sided looks was significantly lower in VPA target individuals, while no difference was observed in partners (LMM: *β* = −7.61 ± 3.42, *t*(58) = −2.23, \**p* = 0.0299 for target individuals; Mann-Whitney U test: *U* = 91, *p* = 0.308 for partners; Fig. 3a). This pattern indicates that the increase in mutual gaze does not stem from greater social motivation in VPA individuals. We then examined gaze responses, defined as orienting toward the partner within 1 second after being looked at. VPA marmosets demonstrated a significantly higher proportion of rapid gaze responses compared with UE controls (LMM: *β* = 8.803 ± 4.187, *t*(58) = 2.102, \**p* = 0.0399; Fig. 3b). This heightened responsiveness was not observed in their partners (Mann-Whitney U test: *U* = 125, *p* = 0.1405; Fig. 3b). Together, these findings indicate a shift toward more reciprocal and less unilateral patterns of gaze coordination in VPA individuals.

**Fig 3.**
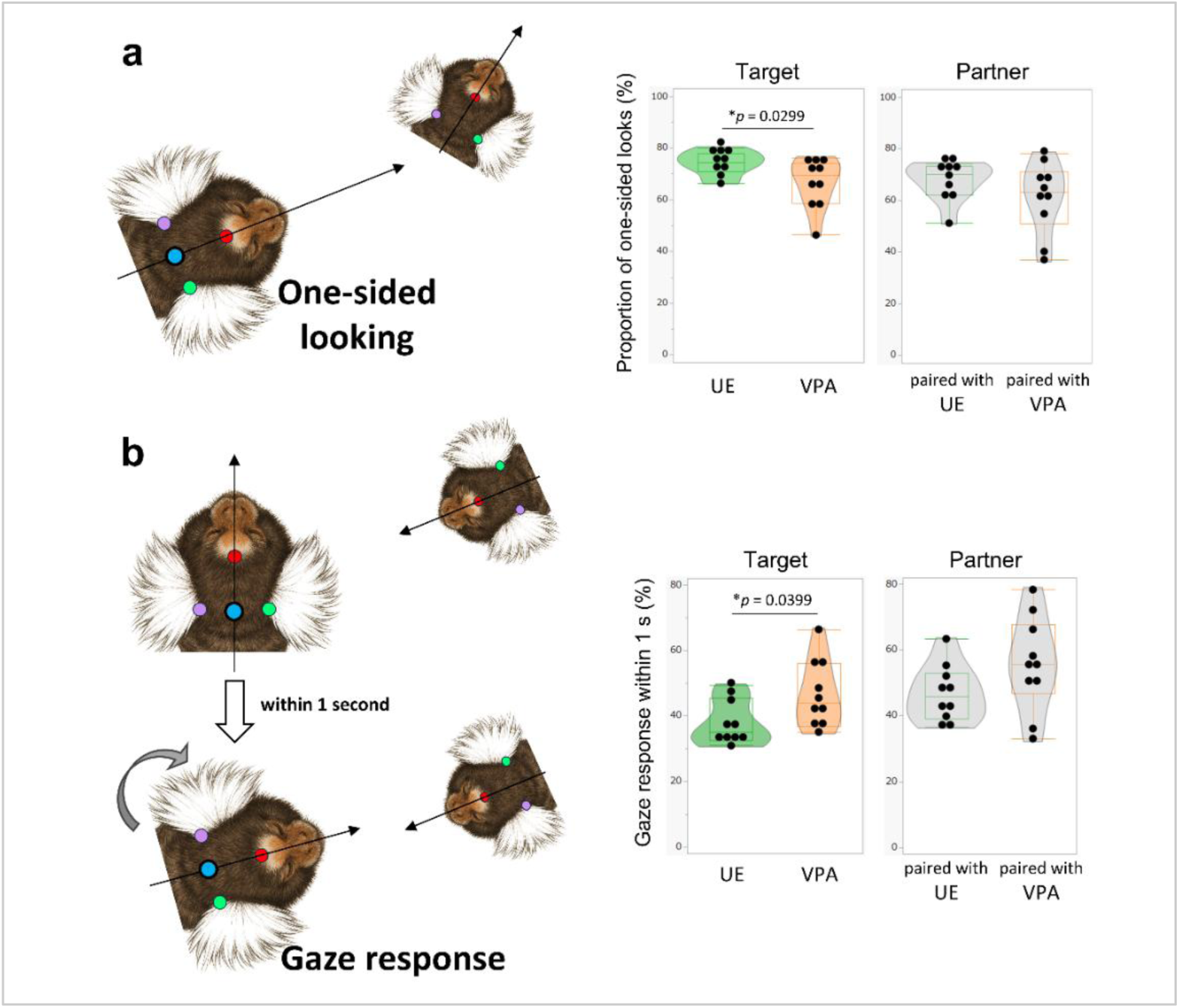
VPA marmosets exhibit fewer one-sided looks and heightened gaze responses. (**a**) Proportion of one-sided looks (%). One-sided looking was defined as the fraction of one individual’s looking events during which the other conspecific did not simultaneously orient toward it. VPA target individuals showed a significantly lower proportion of one-sided looks than UE controls (\**p* = 0.0299), whereas no difference was observed in partners (*p* = 0.308). (**b**) Gaze response within 1 s (%). A gaze response was defined as an event in which one animal oriented toward the other within 1 s after being looked at. VPA target marmosets exhibited significantly higher responsiveness compared with UE controls (\**p* = 0.0399), whereas no difference was observed in partners (*p* = 0.1405). Statistical differences were assessed using a linear mixed model (LMM) for repeated-measures data obtained from the target individuals and the target-partner pairs, and the Mann-Whitney U test for independent measurements from the partners. * = *p* < 0.05.

### Dyad-specific proximity reductions and distance-dependent fragility of mutual gaze in VPA–UE pairs

We next quantified inter-individual distance between the two subjects during the trial. As illustrated by representative trajectories from a UE–UE pair and a VPA–UE pair, target individuals in both conditions often occupied the side of the arena adjacent to the partner (Fig. 4a). To assess spatial occupancy patterns more systematically, we generated two-dimensional heat maps of target individuals’ positions across the entire trial, which confirmed this side preference across both groups (Supplementary Fig. S2). However, despite this shared side preference, the median target-partner distance was significantly shorter in VPA–UE pairs than in UE–UE pairs (LMM: *β* = −85.15 ± 35.11, *t*(18) = −2.426, \**p* = 0.026; Fig. 4b). When distances were computed with respect to a fixed environmental reference (the midpoint of the partner-facing wall; Fig. 4c) rather than to the other animal, no significant group differences were observed for target individuals or for partners (LMM: *β* = −2.06 ± 1.22, *t*(18) = −1.695, *p* = 0.107 for target individuals; Mann-Whitney U test: *U* = 83, *p* = 0.104 for partners; Fig. 4d), indicating that the group effect was specific to dyadic spacing rather than to the absolute positions of either individual. Additionally, gross locomotor measures were comparable: neither total distance moved nor movement area differed significantly for target individuals or for partners (all *p* > 0.05; Supplementary Fig. S2), suggesting that the shorter target-partner distance in VPA–UE pairs is specific to dyadic spacing, rather than reflecting group differences in general locomotor activity or arena-level occupancy. Together, these results indicate that the reduced spacing in VPA–UE pairs reflect a dyad-specific pattern rather than differences in individual positioning or locomotion.

**Fig 4.**
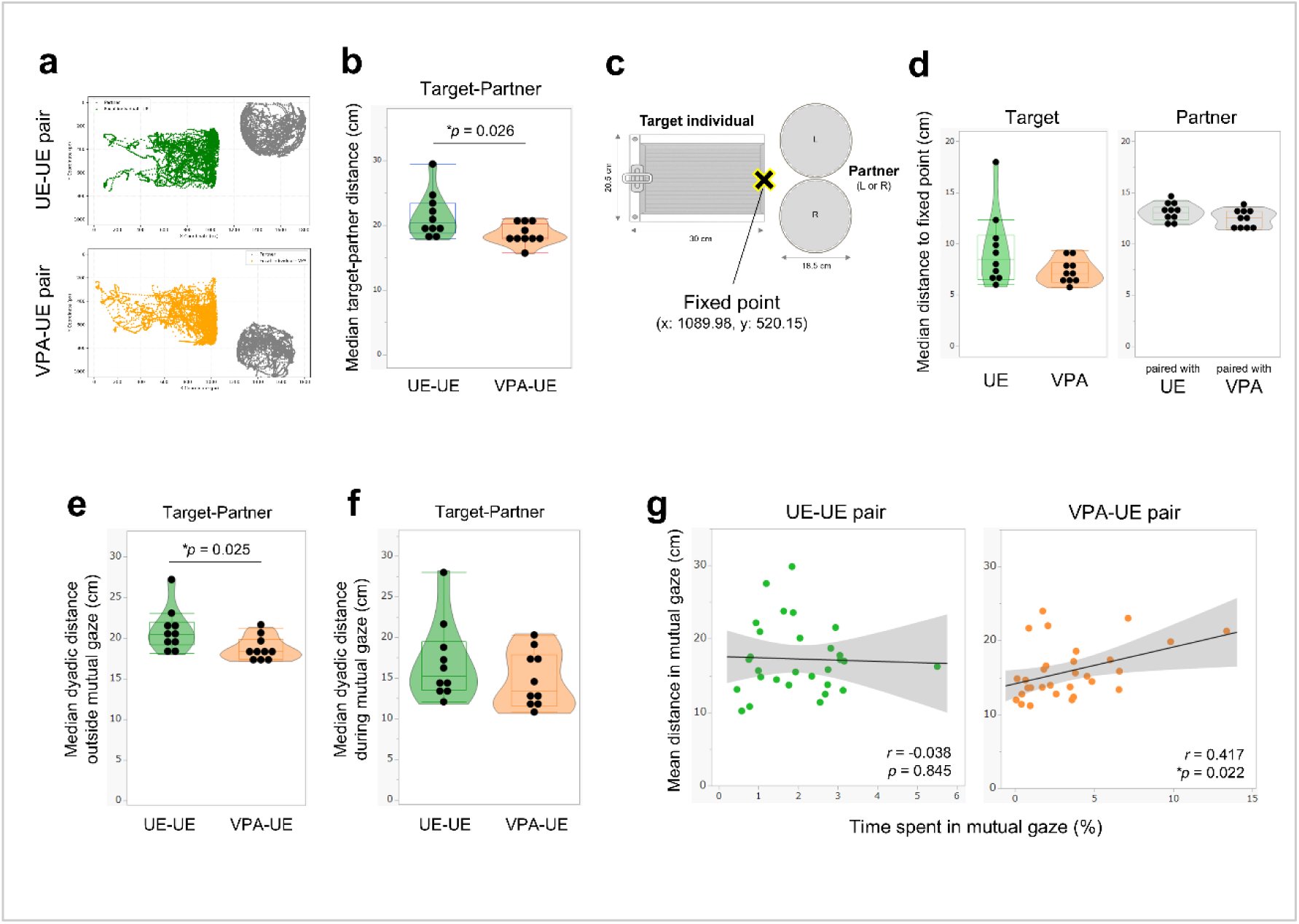
Shorter inter-individual distances and altered coupling with mutual gaze in VPA–UE. (**a**) Overhead view of trajectories from representative trials of a UE–UE pair (top) and a VPA–UE pair (bottom). Trajectories were reconstructed from the frame-by-frame forehead coordinates. (**b**) Violin plots showing median target-partner distance (cm) during the trial. Distances were computed frame by frame from the two individuals’ forehead coordinates and converted to centimeters using arena calibration (1 px = 0.02659 cm). VPA–UE pairs maintained significantly shorter inter-individual distances than UE–UE pairs (\**p* = 0.026). (**c**) Overhead schematic of the apparatus. The fixed point (black X) on the partner-facing wall of the target arena served as a session-invariant reference and was defined as the midpoint of that wall; a single coordinate was obtained by averaging across trials (x = 1089.98, y = 520.15). (**d**) Median distance (cm) from each individual’s forehead to the fixed point, computed separately for the target and the partner. Distances were converted to centimeters using spatial calibration. Although differences did not reach statistical significance, both targets and partners tended to be closer to the fixed point in VPA than in UE (*p* = 0.107 for target individuals; *p* = 0.104 for partners). (**e**) Median target-partner distance outside mutual gaze, computed as in (b) but restricted to frames in which at least one individual’s gaze angle fell outside ±10° relative to the other. VPA–UE pairs maintained shorter distances than UE–UE pairs (\**p* = 0.025). (**f**) Median target-partner distance during mutual gaze, defined as frames in which both individuals’ gaze angles were within ±10° of each other. No significant group difference (*p* = 0.283). (**g**) Relationship between time spent in mutual gaze and mean inter-individual distance during mutual gaze episodes. UE–UE pairs showed no significant correlation (left; *r* = −0.038, *p* = 0.8449), while VPA–UE pairs exhibited a significant positive correlation (right; *r* = 0.417, \**p* = 0.0217). Gray shading indicates 95% confidence intervals. Statistical differences were assessed using a linear mixed model (LMM) for repeated-measures data obtained from the target individuals and the target-partner pairs, and the Mann-Whitney U test for independent measurements from the partners. * = *p* < 0.05.

To test whether this group effect depended on engagement state, we split frames by gaze state. Outside mutual gaze (frames in which at least one individual’s gaze angle fell outside ±10°), the median target-partner distance was shorter in VPA–UE pairs than in UE–UE pairs (LMM: *β* = −2.294 ± 0.937, *t*(18) = −2.448, \**p* = 0.025; Fig. 4e). During mutual gaze (both individuals within ±10°), the median dyadic distance did not differ between groups (LMM: *β* = −1.988 ± 1.789, *t*(15.956) = −1.112, *p* = 0.283; Fig. 4f). Thus, the overall reduction in spacing for VPA–UE pairs was primarily driven by periods without mutual gaze, rather than by the distance maintained during mutual gaze itself.

To investigate the relationship between spatial positioning and gaze behavior, we examined how inter-individual distance during mutual gaze episodes related to the time spent in mutual gaze. In UE–UE pairs, no significant correlation was observed between mean distance during mutual gaze and the percentage of time spent in mutual gaze (*r* = −0.038, *p* = 0.8449; Fig. 4g). In contrast, VPA–UE pairs showed a significant positive correlation between these measures (*r* = 0.417, \**p* = 0.0217; Fig. 4g), indicating that when VPA-UE pairs maintained greater inter-individual distances, they also spent proportionally more time engaged in mutual gaze. Overall, the data suggest a dissociation in VPA–UE pairs between baseline proximity, which is reduced before engagement, and the stability of gaze engagement, which requires an adequate interpersonal distance.

## Discussion

Our findings reveal a specific pattern of dyadic coordination in a non-human primate model of ASD-like behavior, detectable only when the unit of analysis is the pair rather than the individual. VPA–UE dyads displayed a larger proportion and greater frequency of mutual gaze than UE–UE dyads, despite no group differences in overall gaze shifting. VPA-exposed marmosets showed fewer one-sided looks and, when they were looked at by their partner, a markedly higher probability of rapid (within 1 s) gaze responses. Spatially, VPA–UE dyads maintained shorter inter-individual distances than UE–UE dyads; this reduction was specific to dyadic spacing rather than to absolute positioning (no group differences in distance from a fixed environmental reference). When distances were parsed by gaze state, the group difference was evident outside mutual gaze but disappeared during mutual gaze. Moreover, only in VPA–UE pairs did the time spent in mutual gaze increase with greater dyadic distance, whereas UE–UE pairs showed no association. Together, these results indicate a coordination profile that is highly contingent and swiftly responsive to a partner’s cues, yet less tolerant of very close-range engagement. Rather than a blanket “deficit” in social motivation, the pattern suggests context-dependent constraints on how mutual engagement is initiated and sustained.

A growing body of work in human ASD research suggests that gaze avoidance may not primarily reflect reduced social motivation but can, in some contexts, relate to heightened reactivity to direct gaze and close interpersonal proximity. Neuroimaging findings of amygdala and superior temporal responsiveness, together with first-person accounts describing eye contact as intrusive, provide a framework that may be relevant to our observations. In our VPA-exposed marmosets, rapid reciprocal orienting coexisted with reduced stability of mutual engagement at near distances, a combination that could be interpreted as involving proximity-dependent regulatory challenges rather than a simple decrease in social interest. Human hyperscanning studies also indicate that neurotypical partners experience additional perceptual–cognitive demands during interactions with autistic individuals. Consistent with this general pattern, in our dataset, UE partners similarly showed more frequent looking and more dynamic spacing adjustments when paired with VPA animals. These parallels suggest that, in both humans and marmosets, key aspects of social behavior may emerge from dyadic contingencies rather than solely from individual traits.

A central implication is that social interaction differences in animal models of ASD may be mischaracterized if measured only as individual propensities without regard to the partner’s simultaneous state. In our data, VPA target marmosets oriented toward their partners more often (Fig. 2b), but these looks were less likely to occur in isolation (reduced one-sided looking) and more likely to occur as contingent responses (Fig. 3a, b). Partners also altered their behavior: they looked more frequently when paired with a VPA target animal (Fig. 2b), indicating that the dyad’s coordination emerges from both sides. This aligns with interaction-first accounts of social difficulty, which emphasize that measurable differences often reflect dyadic contingencies rather than stable individual traits^11,13,30^. From this perspective, VPA–UE dyads are not “less social”; rather, they appear to occupy a different coordination pattern, in which reciprocal coupling is strong at the sub-second timescale, but close proximity destabilizes sustained engagement. Conceptually, such a regime can generate the seemingly paradoxical combination of more frequent mutual-gaze onsets alongside briefer or more fragile mutual-gaze episodes at short distances, reconciling heightened moment-to-moment responsiveness with reduced tolerance for dense interactional load.

The spatial dynamics observed in our study add another layer of complexity to ASD-related social behavior. VPA–UE pairs maintained significantly shorter inter-individual distances compared to UE–UE controls, suggesting altered preferences for social proximity that may reflect differences in personal space regulation^2,3,31^. Notably, the positive correlation between inter-individual distance and mutual gaze duration specifically in VPA–UE pairs indicates that spatial and gaze coordination are interdependent in VPA-UE dyads. This relationship suggests that when VPA marmosets maintain greater physical distance, they may tend to increase visual engagement, potentially representing an adaptive strategy to optimize social information processing while managing proximity preferences^32,33^. Alternatively, overly close spacing during mutual gaze may introduce aversive stimulation or discomfort in VPA-UE pairs, making sustained visual engagement more difficult; however, the identity of the breaker (the individual who terminates the mutual-gaze episode) did not differ between near- and far-distance episodes. These effects were specific to dyadic spacing: distances to a fixed environmental reference did not differ by group, and gross locomotor metrics, including total distance traveled, were comparable, arguing against general activity differences as an explanation. Taken together, the data suggest that for VPA-UE pairs, a small buffer of inter-individual space may function as a stabilizer that enables the pair to capitalize on intact rapid responsiveness without tipping into destabilizing overload.

Overall, VPA–UE pairs were on average closer yet showed less stable mutual gaze at near distances, suggesting heightened sensitivity to being the target of gaze and early avoidance of near face-to-face contact. In human ASD, difficulties in regulating interpersonal distance are widely reported^3,4,31,34,35^, with seemingly divergent findings of both enlarged interpersonal space and tendencies to approach others too closely. In adults with ASD, stop-distance paradigms often reveal larger preferred or comfortable distances than in controls, accompanied by reduced heart-rate variability, consistent with elevated autonomic arousal and a need for greater space^4,34,36^. At the same time, abnormalities in distance regulation are heterogeneous, and some individuals prefer unusually close spacing^2,3,31^. In our dataset, VPA–UE dyads were on average closer (Fig. 4b), yet mutual gaze was less stable at near distances (Fig. 4g). This combination points to imprecise distance regulation and implies that maintaining an appropriate buffer may be a prerequisite for stable mutual gaze. More broadly, our findings argue against a simple lack of social motivation: VPA animals responded swiftly to partner cues and frequently initiated mutual gaze, but they were less able to sustain it at near distances or for long durations. This profile is consistent with accounts in which social interest is intact but processing of dense social or sensory input is fragile^8,37^. It also accords with reports that ASD individuals modulate eye contact depending on partner and context, as well as with the amygdala hyper-arousal account of gaze avoidance^38,39^.

Although marmosets can be trained for complex tasks, training-based procedures often exclude individuals who do not reach criterion, reducing analyzable samples and risking selection bias. Paradigms that require minimal pre-training and allow natural movement are therefore essential for capturing spontaneous social coordination. The present approach demonstrates how pose-estimation-based tracking can recover fine-grained contingency structure in naturalistic social exchanges. By quantifying head-based gaze proxies frame by frame, we resolved who oriented first, how often one-sided versus reciprocal looks occurred, and how quickly responses followed the partner’s look. These measures extend beyond static “time spent looking” metrics to capture the kinetics and symmetry of social coupling. Notably, these dyad-centric metrics are partner-relative by design. That design choice exposes specificity: the absence of group differences in distance to a fixed wall, together with the presence of robust group differences in distance to the partner, clarifies that the observed effects concern relational organization rather than absolute positioning. More broadly, the pipeline illustrates a path toward standardized, scalable behavioral endpoints that are both ecologically grounded and analytically tractable, providing an important bridge between experimental rigor and the reciprocal nature of real social interaction.

Our study has several limitations that should be acknowledged. First, head orientation is a validated but indirect proxy for eye gaze; brief eye-only deviations cannot be captured without eye tracking^40^. Nonetheless, our ±10° mutual-gaze criterion and consistent geometric definitions reduce ambiguity in classifying interactive looks. Second, the barrier apparatus restricts tactile interaction and imposes a specific spatial affordance; while this control clarifies the visual channel, future work should examine whether the same distance-dependent stabilization holds in fully open encounters or in affiliative contexts that dilute arousal. Third, our paradigm examined initial encounters between unfamiliar individuals, and it remains unclear whether similar patterns would emerge in established relationships or repeated interactions. The marmoset model, while offering valuable social and phylogenetic proximity to humans^18,27,28,41–43^, may not capture all aspects of human ASD presentation, particularly those involving language and complex cognitive processes. Additionally, our analysis focused primarily on target individuals while treating partners as typical controls, potentially overlooking important bidirectional effects that could emerge with more detailed partner characterization. The temporal scope of our observations was also limited to initial interaction periods, and longer-term dynamics or habituation effects remain unexplored.

In sum, the present study reframes ASD-linked social differences in marmosets as an altered pattern of dyadic coordination rather than a simple decrease in engagement. The implications of our findings for intervention design center on dyadic interventions that recognize social difficulties as emerging from interactive dynamics rather than individual deficits alone^11,13,44^. Current interventions often focus on training individuals with ASD to modify their behavior to better match neurotypical expectations^45,46^. Our results suggest that approaches considering both partners’ contributions to successful interaction—including environmental modifications that accommodate altered spatial-visual coordination preferences—may be more effective. For instance, interventions could help neurotypical partners recognize and adapt to different spatial-proximity preferences while supporting individuals with ASD in developing flexible strategies for managing multiple social channels simultaneously. Future research should not only examine how the coordination patterns develop over time and whether they are modifiable, but also integrate concurrent profiling of peripheral physiology and gut microbiome composition and metabolites to test how specific biological states predict or mediate fluctuations in dyadic coordination^47–50^. Such multimodal designs in marmosets could link interactive gaze and spacing dynamics to circulating hormones, inflammatory signals, and microbial diversity or function, generating mechanistic hypotheses for targeted interventions. Understanding the neural mechanisms underlying these dyadic coordination patterns will be crucial for developing targeted, mechanism-based therapeutic approaches that leverage intact social responsiveness while accommodating individual differences in social information processing and regulation.

## Materials and methods

### Animals

Twenty common marmosets (*Callithrix jacchus*) participated as target individuals: 10 prenatal valproic-acid-exposed (VPA) ASD-model animals and 10 unexposed (UE) controls (5 males and 5 females in each group; Supplementary Table S1). Dyads (target and partner) were formed under two conditions, VPA–UE and UE–UE. For each target, three unfamiliar, same-sex UE partners were available; each target completed three sessions, one with a different partner on each occasion. Although the pool of 12 UE partners was reused across targets, every session involved a first-encounter dyad; no target-partner pairing recurred. To minimize carryover from previous encounters, we deliberately scheduled the three sessions months apart (May–July 2023, September–October 2023, and January–February 2024). Partners were pre-screened by experimenters and selected for calm behavior, specifically for remaining composed and not panicking when placed in the apparatus. Target individuals were born and reared in family cages and were then transferred to individual stainless-steel home cages (Natsume Manufacturing Co., Ltd., Tokyo, Japan) at least 2 months before testing. Animals were maintained at 29 ± 2 °C on a 12 h:12 h light:dark cycle (lights on at 07:00 and off at 19:00) with *ad libitum* access to food and water. All animals were habituated to routine handling and readily approached experimenters to obtain food rewards. All experimental animal care procedures were conducted under approved protocols according to the regulations of the National Center of Neurology and Psychiatry (NCNP), Tokyo, Japan (protocol no. 2023033), and the study is reported in accordance with the ARRIVE guidelines.

### Prenatal VPA-exposed ASD marmoset model

Procedures are in accordance with those we have previously reported^26,28^. Briefly, serum progesterone levels in dams were monitored to determine the date of fertilization. VPA was prepared as a 4% solution in 10% glucose and administered intragastrically once daily for seven consecutive days starting on postconception day 60 (200 mg/kg/day). UE dams received no VPA. After birth, regardless of group, infants were housed with both parents in the home cage until weaning.

### Experimental apparatus

All sessions were conducted in a sound-attenuating booth within the experimental room. The booth interior was draped with white fabric to minimize visual distractions, and behavior was monitored on an external display. A rectangular apparatus for target individuals (30.0 cm W × 20.5 cm D × 21.0 cm H; Fig. 1a) and a cylindrical acrylic container for partners (18.5 cm D × 21.0 cm H, wall thickness 0.6 cm; Fig. 1a) were used. The cylindrical case had five circular breathing ports (diameter 0.5 cm) drilled along the lower sidewall in a horizontal row with 10 mm center-to-center spacing. An opaque partition between the two compartments prevented visual contact before trial onset and was operated from outside the booth via a pull cord, allowing trials to commence without exposing the animals to human presence. A video camera (FDR-AX45; Sony Corporation, Tokyo, Japan) was ceiling-mounted inside the sound-attenuating booth and left in place throughout the study so that every trial was recorded from the same fixed viewpoint. The floor was marked with tape to ensure identical placement of the target compartment and partner cylinder across trials. Illumination was provided by two tripod-mounted LED panel lights positioned outside the drape to deliver indirect lighting and avoid reflections in the video; lighting was configured to achieve 2000-2200 lx at the arena surface.

### Procedure

All experiments were conducted in the morning (09:00–12:00). Target individuals voluntarily entered familiar transfer carrier cages from their home cages and were transported to the experimental room. Upon arrival, they were gently guided to voluntarily enter a rectangular testing apparatus. Partners were collected by familiar caregivers from their home cages, voluntarily entered a cylindrical acrylic case designed for the experiment, and were then transported to the experimental room. Transport of the two animals to the room was performed simultaneously by two experimenters to avoid any waiting time for either animal and to minimize differential arousal. Inside the booth, the target compartment and partner cylinder were placed on their pre-marked locations; the partner’s left/right position was counterbalanced across trials according to a preset schedule. Video recording (30 fps) was started once two animals had settled and ended at the conclusion of the 5-min trial. For analysis, we used the 5-min epoch beginning when the partition was opened and both animals were in full, unobstructed view of each other. At the end of each trial, animals were weighed and promptly returned to their home cages.

### Pose estimation (DeepLabCut)

All pose estimation was performed on a Windows 10 Pro (22H2, Intel Core i7-12700KF, 3.60 GHz; 128 GB RAM; NVIDIA GeForce RTX 3090) using Python 3.9.21 (Anaconda 23.3.1), NumPy 1.24.3, and DeepLabCut (DLC) version 2.3.7 with TensorFlow 2.10.0, CUDA 11.2, and cuDNN 8. For network training, we used the first-session videos from each of the 20 target individuals (5 min at 30 fps). Representative frames were selected per video using DLC’s built-in k-means sampling (20 frames/video), yielding 400 labeled frames in total. In every selected frame, six keypoints were annotated: forehead, left ear, and right ear for each of the two animals. The labeled dataset was split such that 95% of frames were used for training and 5% were held out for testing. The network was trained for 150,000 iterations, reaching a final loss of under 0.0015. Pixel errors on the full set were 2.21 px (train) and 6.92 px (test). With a likelihood threshold (p-cutoff) of 0.6 applied, the train error remained 2.21 px and the test error was 6.69 px. The trained model was applied to all experimental videos. Before computing gaze and distance measures, we addressed DLC dropouts on a per-coordinate basis. Short missing runs were linearly interpolated within each time series (bidirectional linear interpolation), whereas longer gaps were preserved as missing to avoid fabricating trajectories. Concretely, sequences of NaNs < 60 frames were filled; sequences ≥ 60 frames (∼2 s at 30 fps) were left as NaN and downstream analyses ignored those frames. The procedure operates column-wise on the 4-level DLC CSV (forehead/ears, x/y, etc.) and saves traceable outputs. After applying the interpolation procedure, we computed detection rates, defined as the percentage of frames in which all required keypoints, and these did not differ between training and held-out test videos for either the target or partner individuals (Supplementary Fig. 2). Agreement at the level of derived measures was also high: the angular difference between gaze vectors computed from human labels versus DLC outputs was sharply centered at 0° for both the target and partner (Supplementary Fig. S3). Together, these checks indicate good generalization and support using DLC-derived keypoints for all downstream analyses.

### Behavioral measures and event definitions

To evaluate whether one subject directed its gaze toward another during the experiment, we employed a geometric method based on body part coordinates obtained via DLC^51^. This approach is particularly justified in New World monkeys such as common marmosets, which rely heavily on head movements for orienting behavior. Previous studies have shown that their eye-in-head movement is limited to approximately ±5° under head-restrained conditions and even further restricted to ±2.5° during free movement, making head orientation a reliable proxy for gaze direction ^15–18^.

The head orientation was estimated by calculating a directional vector from the midpoint of the two ears to the forehead (black arrow; Fig. 1c). This vector was considered to represent the subject’s forward-facing direction. A second vector (blue arrow; Fig. 1c) was defined from the subject’s forehead to the other conspecific’s forehead. This vector served as an estimate of the direction in which the subject would be looking if attending to the other conspecific. To determine whether the subject was looking toward the other animal, we calculated the angular deviation between the head orientation vector and the gaze direction vector. Frames in which this angle fell within 30 degrees were classified as “gaze” frames. An event was labeled “looking” when the inter-vector angle was ≤ 30°, and an event required ≥ 3 consecutive “gaze” frames. To ensure stable azimuth estimates, we required both forehead-left-ear and forehead-right-ear separations to exceed 45 px (1.1966 cm; 1 px = 0.02659 cm); this threshold equals one-half of the empirically measured mean forehead-ear baseline in horizontally oriented images of our subjects and excludes frames in which the head was oriented nearly straight upward or downward in the image, for which 2D gaze-angle estimates are unreliable. We examined the overall distribution of frame-wise gaze angles (Fig. 1d) using circular statistics implemented in Python, pooling all frame-wise angles across individuals and sessions for each pair type (UE–UE and VPA–UE) and role (target vs. partner). Departure from circular uniformity around 0° (direct gaze toward the partner) was evaluated using one-sample V tests against 0°, and Watson’s U² statistics were computed as omnibus measures of deviation from circular uniformity.

Mutual gaze (Fig. 2c, d) was defined more stringently to capture precise face-to-face alignment: both individuals had to satisfy the “gaze” criterion simultaneously with an angular threshold of ≤ 10°. We inspected the onset of every looking event (as defined above) and classified it as a one-sided look (Fig. 3a) if, at that onset frame, the other conspecific did not concurrently satisfy the gaze criterion toward the subject. The one-sided-look ratio was computed per subject and trial as the proportion of looking events that met this condition. Gaze response (Fig. 3b) was recorded if the subject satisfied the looking criterion within a 1-s (≤ 30 frames) window after the prompt onset; otherwise, the prompt was coded as a non-response. Windows truncated by the end of the video were treated as non-responses. At each frame, inter-individual distance (Fig 4b, e, f, g) was computed as the Euclidean distance between forehead landmarks which is the square root of the sum of the squared horizontal and vertical separations between the two foreheads and then converted from pixels to centimeters (1 px = 0.02659 cm). We also computed the distance from each individual’s forehead to a fixed point in the target arena (Fig. 4c, d) using the same Euclidean formula as for inter-individual distance, substituting the fixed-point coordinates.

### Statistical Analysis

To compare group differences in behavioral indices measured from the subject individual, linear mixed-effects models (LMMs) were applied separately to each behavioral variable. The variables analyzed included median distance, frequency of gaze events, gaze duration, and other behavioral indices derived from the subject individual. In each model, group (UE vs. VPA) was treated as a fixed effect, and subject identity was included as a random intercept to account for repeated measurements. Since each subject interacted with multiple different partner individuals across sessions, this modeling approach allowed us to retain trial-level variability while appropriately accounting for within-subject dependence. The models were fitted using the *lmerTest* package (version 3.1.3) in R (version 4.3.3). Model assumptions regarding the normality and homoscedasticity of residuals were evaluated using diagnostic plots and the Shapiro-Wilk test. Fixed effects were assessed using Satterthwaite’s approximation for degrees of freedom. For the partner individual, who differed in each session and was not subject to repeated measurements, group comparisons of behavioral indices were conducted using either parametric or nonparametric tests, depending on the distribution of the data. Specifically, normality and homogeneity of variance were assessed, and when parametric assumptions were not met, nonparametric Mann-Whitney U tests were performed. These tests were conducted in JMP Pro (version 17.2.0; SAS Institute Inc., Cary, NC, USA).

## Data availability

The datasets used and/or analyzed during the current study are available from the corresponding author on reasonable request.

## Acknowledgments

We thank Dr. Atsushi Senjyu from Hamamatsu University School of Medicine for constructive feedback. We are grateful to Dr. Jun Noguchi, Dr. Satoshi Watanabe and Dr. Kazuhiko Yamaguchi from NCNP for their helpful discussions. We also thank to Dr. Naomi Hasegawa, Akiko Tsuchiya, Ryoichi Saito, and other members of the NCNP primate facility for their invaluable support with experiments and animal care. Finally, we extend special thanks to all the subject marmosets.

## Authors information

### Contributions

M.N. designed the study, conducted experiments, analyzed the data, and wrote the manuscript; Tomoko S. conducted experiments and analyzed the data; T.K. designed and conducted experiments; K.S. conducted experiments; A.M.H. and S.S. carried out data collection; K.M. contributed to study design and provided comments on the manuscript; N.K. provided comments on the manuscript; Tsuyoshi S. supervised the project, acquired funding, and provided comments on the manuscript. N.I. designed the study, acquired funding, supervised the project, and wrote the manuscript. All authors approved the final version of the manuscript and agree to be accountable for all aspects of the work.

