## Supplemental Table S1, Figure S1-S3 for "Heightened gaze responsiveness and mutual gaze instability at close distances characterize dyadic coordination in a marmoset model of autism"

**Supplementary Table S1.** Animal information. Twenty common marmosets (*Callithrix jacchus*) participated as target individuals (10 VPA, 10 UE). Twelve UE marmosets were used as partners; some individuals participated in sessions with more than one target. Animals with the same symbol in the "Siblings" column are littermates. Age (year) and body weight (g) are per-subject means across the experimental sessions. Body weight at testing did not differ between target groups (UE: 382.7 ± 42.5 g; VPA: 385.0 ± 27.5 g; Welch's $t(15.4) = 0.1427$, $p = 0.888$).

| *Group* | *Role* | *Animal ID* | *Age (year)* | *Siblings* | *Sex* | *Weight (g)* |
|---|---|---|---|---|---|---|
| UE | Target | 16086 | 7.2 | | Female | 364.2 |
| UE | Target | 16089 | 7.2 | & | Male | 443.0 |
| UE | Target | 16090 | 7.2 | & | Male | 362.4 |
| UE | Target | 16091 | 7.2 | | Female | 410.6 |
| UE | Target | 16104 | 7.1 | | Male | 344.7 |
| UE | Target | 17013 | 6.7 | | Male | 368.8 |
| UE | Target | 17015 | 6.7 | | Male | 466.7 |
| UE | Target | 17104 | 5.9 | | Female | 346.0 |
| UE | Target | 20156 | 3.1 | = | Female | 355.6 |
| UE | Target | 20157 | 3.1 | = | Female | 365.5 |
| VPA | Target | 14002 | 9.5 | | Female | 380.4 |
| VPA | Target | 16018 | 7.3 | | Male | 396.8 |
| VPA | Target | 16109 | 7.1 | | Female | 400.0 |
| VPA | Target | 16153 | 6.9 | | Female | 349.7 |
| VPA | Target | 17024 | 6.5 | # | Male | 374.9 |
| VPA | Target | 17026 | 6.5 | # | Male | 448.1 |
| VPA | Target | 19061 | 4.2 | $ | Male | 359.9 |
| VPA | Target | 19062 | 4.2 | $ | Female | 384.4 |
| VPA | Target | 21070 | 2.6 | + | Female | 391.2 |
| VPA | Target | 21071 | 2.6 | + | Male | 364.8 |
| UE | Partner | 13024 | 10.4 | | Male | 345.4 |
| UE | Partner | 16045 | 7.2 | | Female | 348.5 |
| UE | Partner | 16077 | 7.2 | | Male | 369.1 |
| UE | Partner | 16112 | 7.1 | | Female | 360.7 |
| UE | Partner | 16137 | 7 | | Male | 380.4 |
| UE | Partner | 17086 | 6.3 | | Female | 409.5 |
| UE | Partner | 17149 | 5.8 | | Female | 372.6 |

| | | | | | | |
|---|---|---|---|---|---|---|
| UE | Partner | 18014 | 5.5 | | Female | 440.2 |
| UE | Partner | 18027 | 5.5 | | Female | 348.7 |
| UE | Partner | 19025 | 4.6 | | Male | 345.5 |
| UE | Partner | 19137 | 4.2 | | Female | 451.9 |
| UE | Partner | 20145 | 2.9 | | Male | 362.5 |

**Supplementary Video S1.** Example of dyadic interaction with DLC-based gaze vectors. Representative 10-s excerpt from a first-encounter session of a dyad; the target individual is shown on the left and the partner on the right. The positions of the forehead and both ears detected by DLC are overlaid as keypoints, and head-orientation gaze vectors are shown as arrows for both animals. The vectors are displayed only for frames that passed the azimuth validity criterion described in the Methods; arrows are therefore absent when the head is oriented substantially upward or downward in the image, for which 2D gaze-angle estimates are unreliable. The video is shown in real time (30 fps).

**Supplementary Figure S1.**

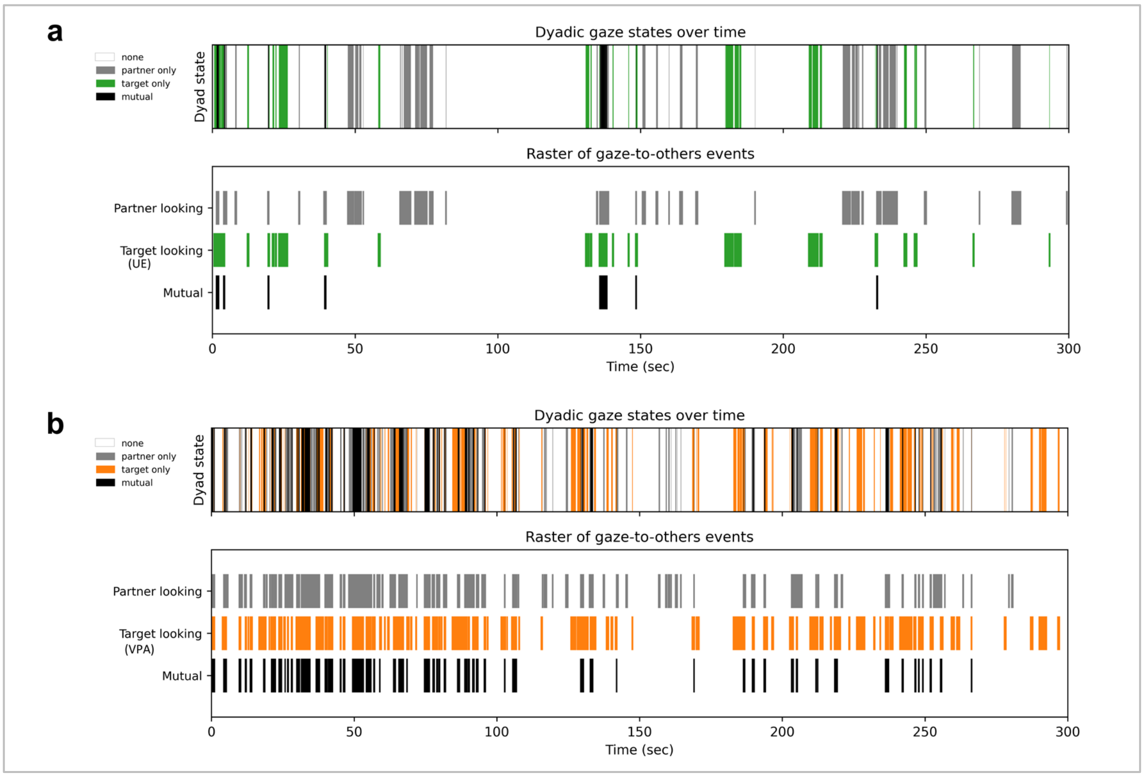


**Supplementary Figure S1.** Representative dyadic gaze timelines. (**a**) UE–UE pair and (**b**) VPA–UE pair. Each panel shows representative dyadic-gaze data from a single dyad selected as a typical example of the group. The upper plot displays the frame-by-frame dyadic state over time (0-300 s): white, neither individual looking at the other conspecifics (“none”); gray, only the partner individual looking at the target (“partner only”); green (UE target)/orange (VPA target), only the target individual looking at the partner (“target only”); black, simultaneous looking by both individuals (“mutual”). The lower plot shows a raster of gaze-to-others events for the same session: from top to bottom, partner looking, target looking, and mutual gaze. Mutual gaze (black) was defined using the same criteria as in the main analyses (gaze vectors within ±10°, distance and tracking-quality thresholds satisfied, and ≥3 consecutive frames).

**Supplementary Figure S2.**

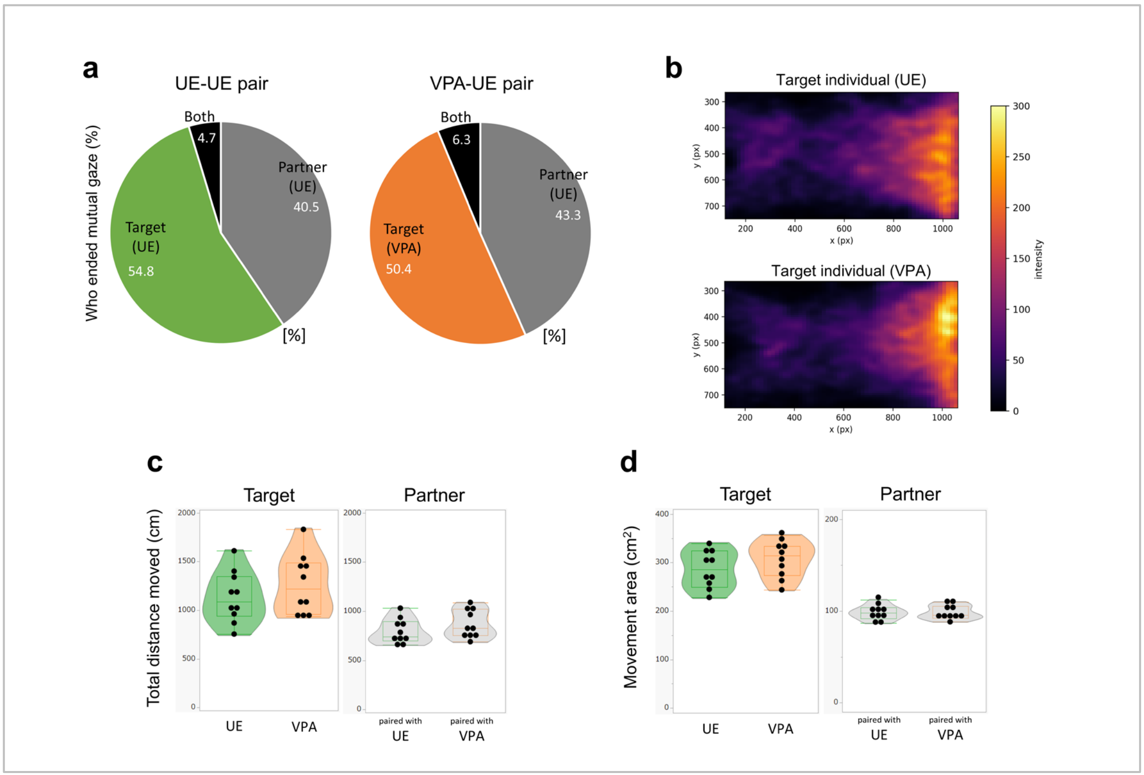


**Supplementary Figure S2.** Control analyses of mutual gaze termination and locomotor activity. (**a**) Proportion of mutual-gaze events attributed to the target vs. partner as the terminator of the event; "Both" indicates simultaneous termination within the same video frame. (**b**) Two-dimensional occupancy heat maps for target individuals, aggregated across sessions. (**c**) Total distance moved (cm) for targets and for partners. No between-group differences were detected (LMM: $\beta = 136.03 \pm 126.78$, $t(18) = 1.073$, $p = 0.297$ for target individuals, Mann-Whitney U test: $U = 123$, $p = 0.186$ for partners). (**d**) Movement area ($cm^2$) for targets and partners. No between-group differences were detected (LMM: $\beta = 20.66 \pm 16.95$, $t(18) = 1.219$, $p = 0.238$ for target individuals, Mann-Whitney U test: $U = 103$, $p = 0.910$ for partners). Dots show per-subject means across the three experimental sessions; violins depict kernel density with the median indicated.

**Supplementary Figure S3.**

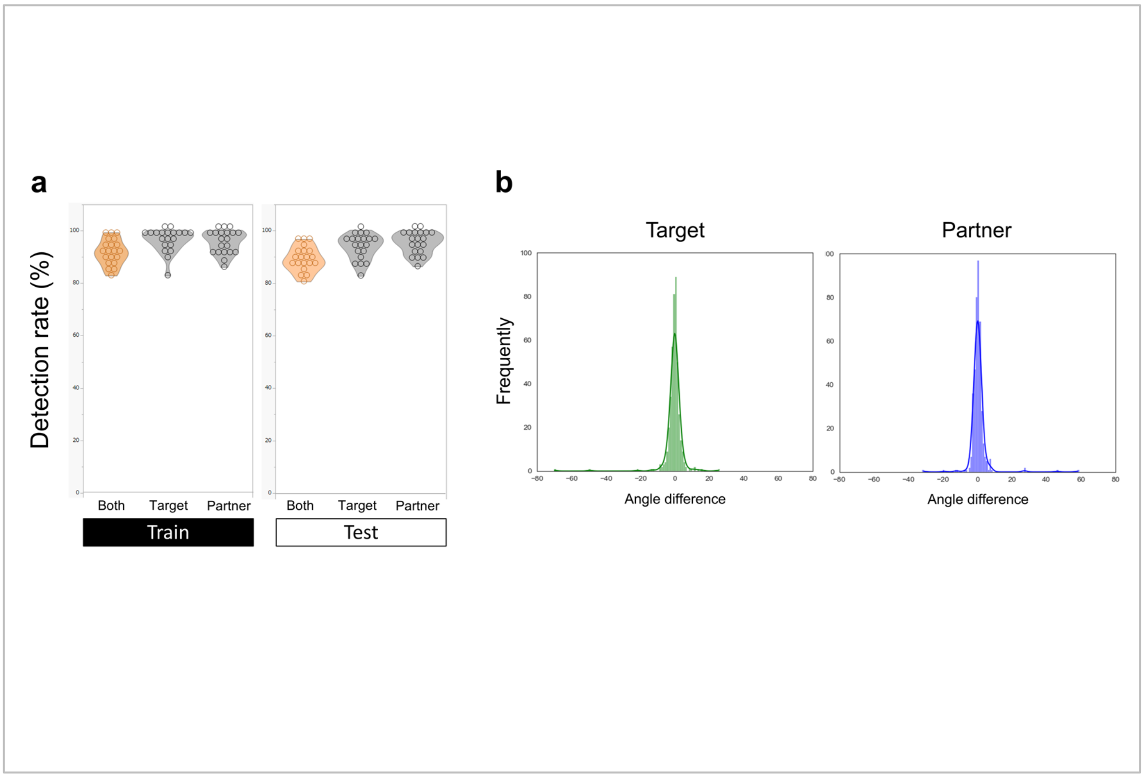


**Supplementary Figure S3.** DLC keypoint detection and human–model agreement. (**a**) Detection rate (%) of the three DLC keypoints (forehead and both ears) used for gaze computation, shown for frames in which both animals were valid ("Both") and for each individual (Target, Partner), split by videos used for Train (left) vs Test (right). Detection rates were near ceiling across conditions, with no systematic difference between train and test sets. Dots indicate per-subject values; violins show the distribution with median lines. (**b**) Agreement between human annotations and DLC estimates of head-orientation gaze: distributions of frame-wise signed angle differences (degrees) for the target (left, green) and partner (right, purple). Both distributions are narrowly centered around 0°, indicating close correspondence between manual and DLC-derived vectors. All validity filters match those described in Methods.
